# AAV2-Retro-Mediated Gene Transfer Selectively Targets Outer Retinal Cells Following Intravitreal Injection

**DOI:** 10.64898/2026.03.10.710806

**Authors:** Chaimaa Kinane, Moxa Panchal, Pantelis Tsoulfas, Venu Talla, Kevin K. Park

**Affiliations:** Department of Ophthalmology, University of Texas Southwestern Medical Center; Department of Neuroscience, University of Texas Southwestern Medical Center; Peter O’Donnell Jr. Brain Institute, University of Texas Southwestern Medical Center; Department of Neurosurgery, The Miami Project to Cure Paralysis, University of Miami Miller School of Medicine

## Abstract

**Purpose:** To characterize the cellular tropism and temporal dynamics of adeno-associated virus 2 (AAV2)-retro–mediated gene delivery in the adult mouse retina following intravitreal injection.

**Methods:** Adult C57BL/6J mice received single or sequential intravitreal injections of AAV2-retro carrying the mGreenLantern (mGL) reporter gene. Retinas were collected at 1-, 3-, and 14-days post-injection (dpi) and processed for immunofluorescence analysis. Transduced cell types were identified using cell-type markers, including cone arrestin, RBPMS, and AP-2α. The number and distribution of mGL-positive cells were quantified on whole retinas or retinal cross-sections to assess transduction efficiency, specificity, and spatial coverage.

**Results:** Reporter expression was detected in the outer retina at 1 dpi and increased markedly at 3 and 14 dpi. AAV2-retro demonstrated strong tropism for photoreceptors and retinal pigment epithelium (RPE), with robust labeling of both rods and cones. In contrast to the robust outer retinal expression, transduction in the inner nuclear layers was limited to a few retinal ganglion and amacrine cells, reflecting strong cell-type specificity. Reporter expression was distributed widely across the retina, exceeding the localized pattern typically observed following subretinal delivery with conventional AAV2 vectors. Sequential injections further increased reporter expression and spatial coverage compared with single injections.

**Conclusions:** AAV2-retro enables efficient, outer retina–specific gene delivery following intravitreal administration. This approach overcomes the limitations of traditional intravitreal gene transfer and provides a minimally invasive alternative to subretinal injection. AAV2-retro– mediated transduction may facilitate preclinical studies of retinal degeneration and support the development of gene therapies aimed at preserving photoreceptors and RPE function.

## INTRODUCTION

Inherited and acquired retinal degenerative diseases are a leading cause of irreversible blindness worldwide, primarily due to the loss of photoreceptors or dysfunction of the supporting retinal pigment epithelium (RPE).^1–3^ Gene therapies targeting these cells offer a promising avenue for restoring vision or halting disease progression.^4–7^ Recombinant adeno-associated virus (rAAV) vectors have emerged as a key platform in this field, demonstrating sustained gene expression, low immunogenicity, and a strong safety profile in both ocular and non-ocular applications. In particular, rAAV-mediated gene transfer to photoreceptors and RPE has shown significant therapeutic potential in preclinical models and human clinical trials.^7–11^ Currently, a widely used method for delivering AAV vectors to photoreceptors is subretinal injections, which place the vector in direct contact with the outer retina.^12–14^ This approach is exemplified by the FDA-approved gene therapy voretigene neparvovec (Luxturna), which uses AAV serotype 2 (AAV2) to restore RPE65 function in patients with inherited retinal dystrophy.^15^ However, subretinal delivery requires a complex surgical procedure involving retinal detachment and the creation of a transient subretinal bleb.^16^ This is not only technically demanding but also poses risks to an already fragile retina, particularly in late-stage disease when retinal structure is severely compromised.^17,18^ Moreover, subretinal injections typically result in localized gene expression confined to the area of the bleb, as conventional AAV capsids are limited in their ability to diffuse laterally through retinal tissue.^4,19^ Consequently, only a portion of the retina is transduced, which may be insufficient to halt disease progression or restore functional vision, especially in conditions that require widespread retinal coverage. These limitations underscore a critical need for alternative delivery strategies that are less invasive and more broadly effective.

Intravitreal injection, by contrast, is a well-established, minimally invasive technique commonly used for the delivery of pharmacological agents in clinical ophthalmology. Its accessibility and repeatability make it an attractive route for ocular gene therapy. However, the inner limiting membrane (ILM) and other physical barriers of the inner retina hinder AAV access to deeper retinal layers, including photoreceptors and the RPE, limiting the utility of traditional AAV serotypes when delivered intravitreally.^20–23^

Recent studies have identified engineered AAV capsids with enhanced ability to traverse retinal barriers and reach the outer retina from the vitreous cavity.^23–26^ Others have identified promoters which confer highly selective gene expression in the outer retina.^27^ These novel AAV variants and promoters are under active investigation and hold significant promise, though their efficiency, safety, and translational readiness for broad clinical application are being established through ongoing studies.

AAV2-retro is an engineered AAV capsid variant derived from AAV2, specifically developed to enhance retrograde transport in neurons. Originally described by Tervo et al. in 2016, AAV2-retro enables robust gene delivery to projection neurons by traveling against the direction of synaptic transmission^28^. In this study, we initially sought to determine whether AAV2-retro, when delivered via intravitreal injection, exhibits a preferential pattern of transduction in retinal neurons that reflects retrograde transport through the visual circuitry and, if so, identify which neuronal cell types in the retina are most efficiently transduced. Here we report that intravitreal delivery of AAV2-retro results in robust transduction of photoreceptors and RPE cells within one day, demonstrating a rapid onset and striking cell-type specificity for these cell types in the adult mouse retina.

These findings highlight the unexpected capability of AAV2-retro to bypass inner retinal barriers. The capacity to target outer retinal cells non-invasively in adult animals offers major advantages for preclinical studies, enabling more accessible, reproducible, and less damaging gene delivery in animal models of retinal degeneration and repair.

## MATERIAL AND METHODS

### Ethical review

All animal procedures were conducted in compliance with protocols approved by the Institutional Animal Care and Use Committee (IACUC) at University of Texas Southwestern Medical Center (UTSW) and in accordance with the Association for Research in Vision and Ophthalmology Statement for the Use of Animals in Ophthalmic and Vision Research.

### Animal

Male and female C57BL/6J mice (6–8 weeks old; Jackson Laboratory, stock no. 000664) were used for this study. Postnatal day 3 (P3) pups, regardless of sex, were used for the postnatal injection experiment. Mice were housed in individually ventilated cages under a 12 h light/Dark cycle. Food and water were provided *ad libitum*, and nesting material and shelters were added to all mice cages for environment enrichment.

### Intravitreal injection

Mice were anesthetized using Isoflurane (3% for induction, and 1.5-2% for maintenance) vaporized in oxygen. Intravitreal injections were performed using pulled glass micropipettes under a stereomicroscope as described previously.^29,30^ A total volume of 2 μl of viral vector was injected into the vitreous chamber of the right or left eye through the sclera just posterior to the limbus.

Mice received intravitreal injections of either AAV2-retro-CMV-H2B-mGL, AAV2-CMV-H2B- mGL or AAV MNM008-CAGIG-H2B-V5-mGL (approximately 1.8 x 10^10^ vg/eye). The original titers of these AAVs were AAV2-retro-CMV-H2B-mGL (3.46 x 10^13^ vg/ml), AAV2-CMV-H2B- mGL (0.9 x 10^13^ vg/ml), and AAV MNM008-CAGIG-H2B-V5-mGL (7.82 x 10^13^ vg/ml). Buprenorphine SR was administered subcutaneously as post-operative analgesia.

For the postnatal injection study, intravitreal injections were performed at P3. Pups were anesthetized by hypothermia on ice, and viral vectors were delivered into the vitreous chamber using pulled glass micropipettes under a stereomicroscope. After injection, pups were allowed to recover on a warming pad before being returned to the dam.

### AAV vectors and virus production

AAVs were produced at the UTSW Ophthalmology Gene Editing and Viral Production Core or at the University of Miami Viral Vector Core. The reporter plasmid pAAV.CMV.H2B.mGL.WPRE contains AAV2 ITRs, a CMV enhancer/promoter, a chimeric intron, the mGreenLantern (mGL) reporter fused to histone H2B, an hGH polyadenylation signal, and a WPRE element (Supplemental Figure S1). AAVs were produced with two different capsids the AAV2-retro (rAAV2-retro helper was a gift from Alla Karpova & David Schaffer (Addgene plasmid # 81070; RRID:Addgene_81070) and MNM008.^11^ The MNM008 capsid was constructed on the AAV2- retro backbone plasmid by substituting the sequence between N587 and R602 of the AAV2-retro VP1 capsid gene with the amino acid sequence SFTSPLHKNENTVS. The viruses were purified following previously described protocols^31^, and viral titers were quantified using either qPCR or digital PCR.

### Tissue processing

At designated time points (1 day, 3 days, and 2 weeks post-injections), mice were humanely euthanized in accordance with the American Veterinary Medical Association guidelines by CO₂ inhalation followed by a confirmatory secondary method. Once the absence of reflexes was confirmed, animals were transcardially perfused with ice-cold phosphate-buffered saline (PBS, 1X), followed by 4% paraformaldehyde (PFA) in PBS. Eyes were then enucleated and post-fixed in 4% PFA at room temperature (RT) for 2-3 hours before further processing. For flat mount retinal preparation, retinas were carefully dissected out from the eye cup and were washed three times in PBS. Four radial cuts were made at the ora serrata to create a cloverleaf shape before mounting.

For cryosections, eyes were cryoprotected by immersion in 30% sucrose at 4 °C overnight. Tissues were embedded in optimal cutting temperature (OCT) compound using cryomolds and sectioned at a thickness of 10 μm on a Leica CM3050S cryostat.

### Immunohistochemistry

To perform the immunofluorescent staining, samples were blocked with PBS containing 10% normal goat serum (NGS) and 0.3% Triton X-100 (0.3% PBST) for 1 hr hour at room temperature (RT). Samples were then incubated with anti-RBPMS antibody (1:2,000; Custom made at ProSci), anti-Cone arrestin (1:200; cat# AB15282; Sigma Aldrich), and anti- AP-2α antibody (1:50; cat# 3B5; DSHB) that were diluted in PBST blocking solution and incubated at 4 °C overnight. After 3 washes (10 minutes each) in PBST, sections were incubated with anti-guinea pig Alexa Fluor 594 (1:500; cat# 106-585-003; Jackson ImmunoResearch Laboratories), anti-rabbit Alexa Fluor 647 (1:500; cat# ab150079; Abcam), and anti-Mouse Alexa Fluor 647 (1:500; cat# ab150115; Abcam) for 2 hours at RT. Samples are rinsed 3 times with PBST and then mounted onto a glass coverslip using Fluomount-G Mounting Medium.

### Imaging and quantification

Retinal sections were imaged using a confocal microscope (Zeiss inverted 780). Images were acquired using 20X and 40X objectives. Sequential acquisition was performed to avoid spectral overlapping between fluorophores. Z-stacked images were acquired, and maximum intensity projections were generated using Fiji (Image J). Layer identification and qualitative assessment were performed based on DAPI staining and anatomical landmarks. For quantification, single-plane images were used, and maximum intensity projections were generated for visualization.

Cell quantification was performed on at least two fixed rectangular region of interest (ROI) measuring approximatively 59 x 198 um, applied consistently across sections and animals. At least three animals were used for quantification.

Data are presented as mean ± SEM and analyzed using GraphPad Prism. A *P* value < 0.05 was considered significant (* *P* < 0.05, ** *P* < 0.01, *** *P* < 0.001, and **** *P* < 0.0001).

## RESULTS

### Intravitreal injection of AAV2-retro leads to rapid and preferential transduction of photoreceptors and RPE cells

To evaluate the transduction profile of AAV2-retro following intravitreal injections, we examined retinal tissue at 1-, 3-, and 14-days dpi in adult male mice. At 1 dpi, the reporter mGL expression was already detectable, predominantly localized to the outer retina (Figs. 1A, 1B), suggesting early onset of transgene expression in photoreceptors and RPE cells. By 3 dpi, transduction intensity increased markedly, with a broader distribution across the photoreceptor layer and consistent labeling of RPE cells (Figs. 1C, 1D). Fluorescence signal intensity and cellular coverage were both elevated compared to the 1-day time point, indicating active viral entry and transcription. At 3 dpi, only a few GFP^+^ cells were seen in the ganglion cell layer (GCL) (Fig. 1D).

**Figure 1.**
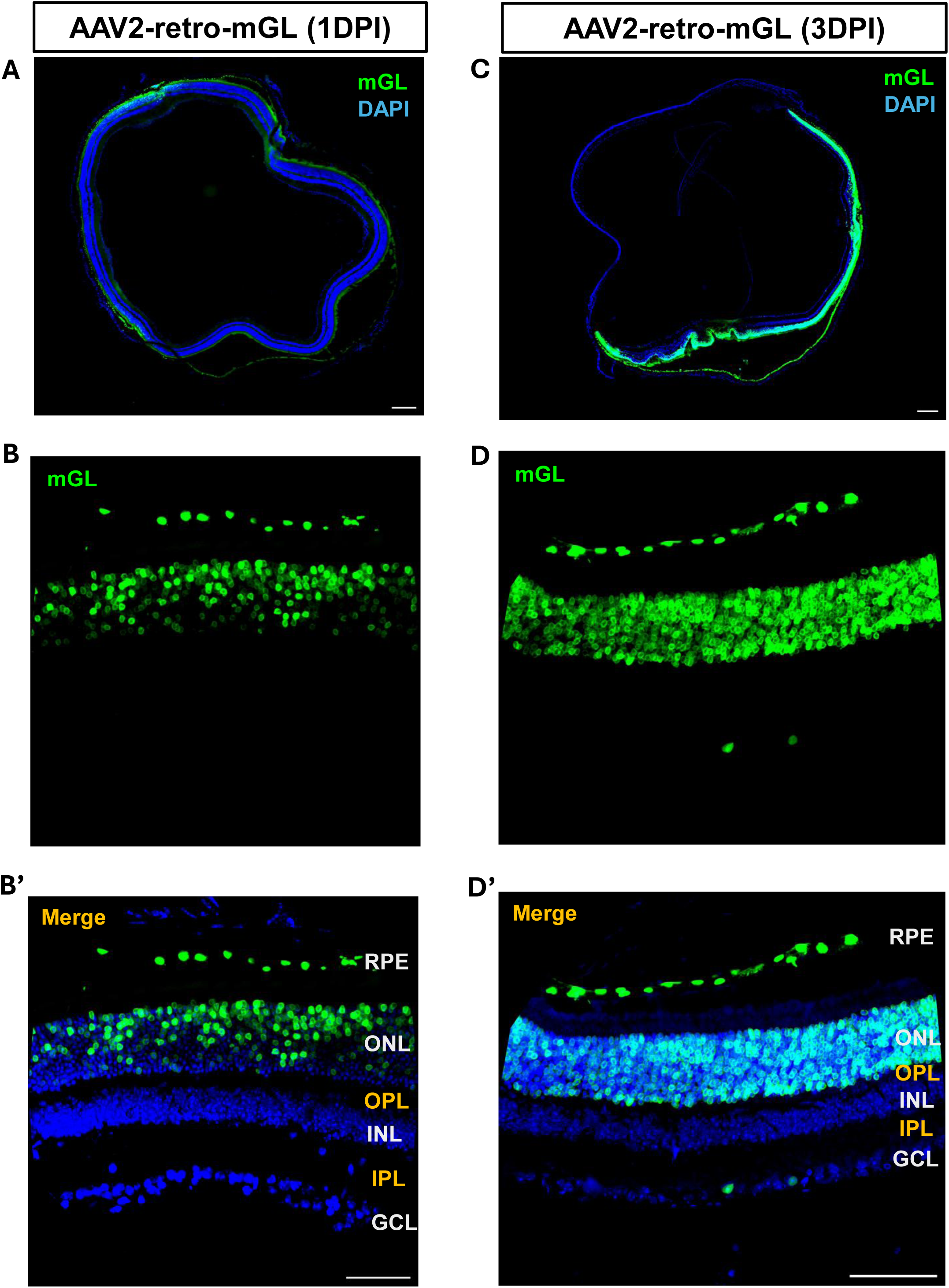
AAV2-retro preferentially transduces photoreceptors and RPE 1 and 3 days after intravitreal injection. Stitched 10X images of retinal cross-sections showing mGL expression at 1 day post injection (dpi) **(A)** and 3dpi **(C)**. DAPI (blue) labels nuclei; green indicates mGL reporter expression. **(B and D)** Higher-magnification images of representative retinal regions showing mGL expression at 1 dpi (B) and 3 dpi (D). **(B’ and D’)** Merged images corresponding to panels B and D, illustrating mGL and DAPI expression across retinal layers. Retinal layers are indicated: RPE, ONL, OPL, INL, IPL, and GCL. *Scale bars*: 200 µm (A, C); 50 µm (B’, D’).

At 14 dpi, a robust GFP expression was observed throughout the photoreceptor layer, with strong labeling of RPE cells. (Figs. 2A, 2B) Minimal signal was observed in the GCL and the inner nuclear layer (INL) (Fig. 2B), confirming the tropism of AAV2-retro for outer retinal cell types under these conditions. These results demonstrate that intravitreal delivery of AAV2-retro in adult mice leads to rapid, preferential, and progressively enhanced transduction of photoreceptors and RPE cells within two weeks post-injection.

**Figure 2.**
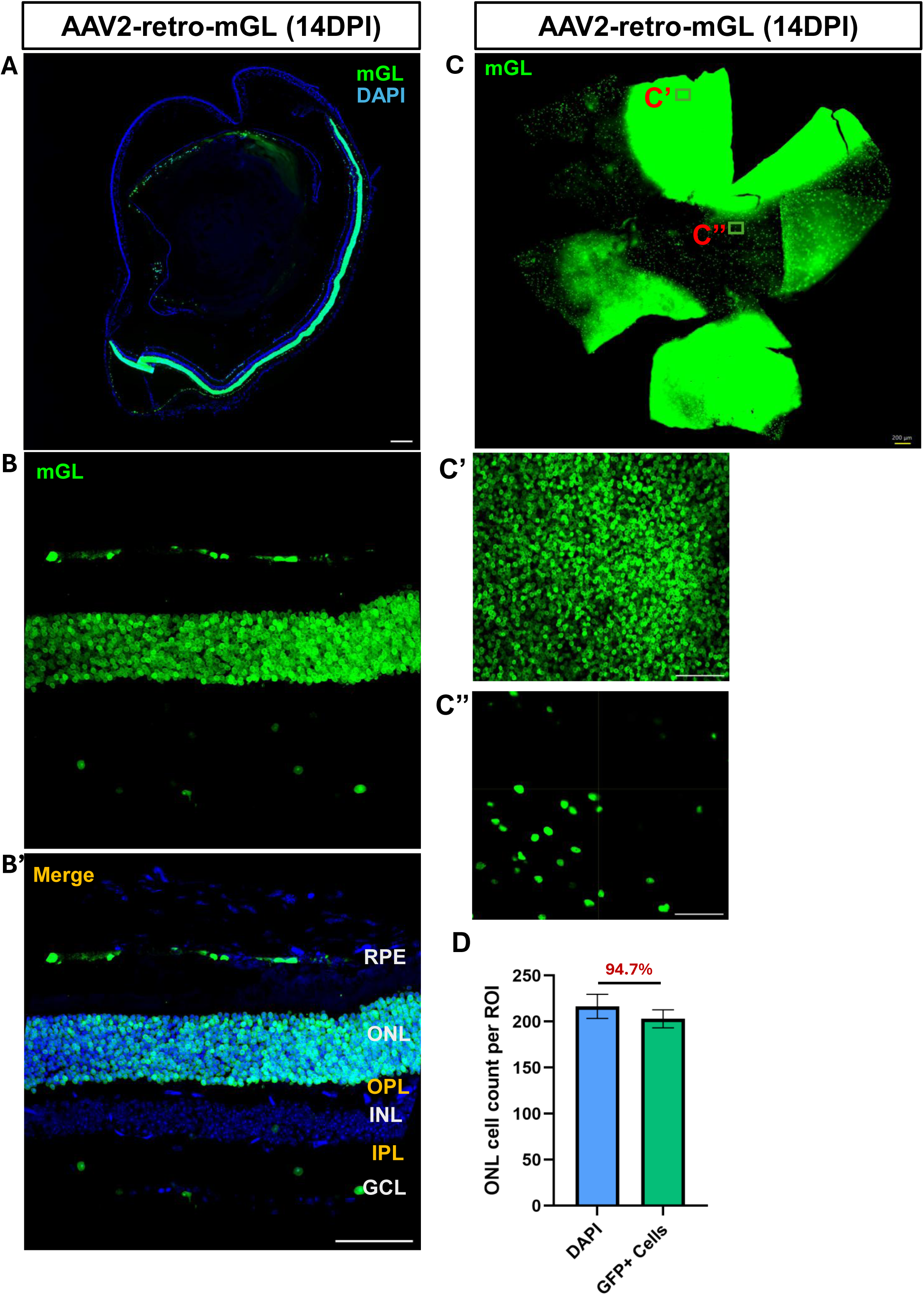
AAV2-retro maintains selective transduction in the outer layers 14 days after intravitreal injection. (A) Stitched 10X images of retinal cross-sections showing mGL expression at 14 dpi. (B) Higher-magnification view of a representative retinal region with mGL expression. (B’) Merged image from (B) highlighting layer-specific distribution (RPE, ONL, OPL, INL, IPL, and GCL), mGL, and DAPI across the layers. (C) Retinal flat-mount at 14 dpi after sequential injections, showing widespread mGL expression. (C’ and C’’) Higher magnification images of the boxed areas in (C). Scale bars: 200 µm (A, C); 50 µm (B’, C’, C’’). (D) Quantification of mGL- positive cells within the regions of high transduction in the ONL (i.e., region of interest, ROI). The percentage shown indicates the proportion of transduced cells among total DAPI-positive ONL cells. N= 3 animals. *Scale bars*: 200 µm (A, C); 50 µm (B’, D’).

Normally, AAV-mediated transduction following a single injection is spatially concentrated, with robust expression near the presumed injection site. To assess whether sequential AAV2-retro delivery improves transduction efficiency in wider area, adult mice received two intravitreal injections spaced three days apart. Retinal tissues were analyzed 14 days after the initial injection. Compared to a single injection, sequential dosing resulted in markedly increased reporter expression across the retina, with broader distribution and higher intensity in both photoreceptors and RPE cells (Fig. 2C). In some retinas, reporter expression extended to nearly the entire retina, and this widespread coverage was already observed by 3dpi (data not shown). In regions with strong mGL expression, over 90% of DAPI-stained cells in the photoreceptor layer were positive for mGL 14 days post-injection (Fig. 2D).

### AAV2-retro exhibits similar photoreceptors and RPE transduction in female mice

So far, we used male mice. To assess whether the AAV2-retro transduction pattern observed in males was conserved across sex, intravitreal AAV2-retro injections were performed in adult female mice. Analysis of retinal cross-sections revealed robust reporter expression predominantly localized in the RPE and ONL, with strong photoreceptors and RPE transduction. This ONL- transduction pattern was evident at both 3 and 14-dpi and closely resembled the distribution observed in male retinas (Fig. 3). Minimal reporter expression was detected in the INL and GCL. These findings were consistently observed in three female animals, indicating that AAV2-retro– mediated transduction of RPE and outer retinal cell types is comparable between sexes under these conditions.

**Figure 3.**
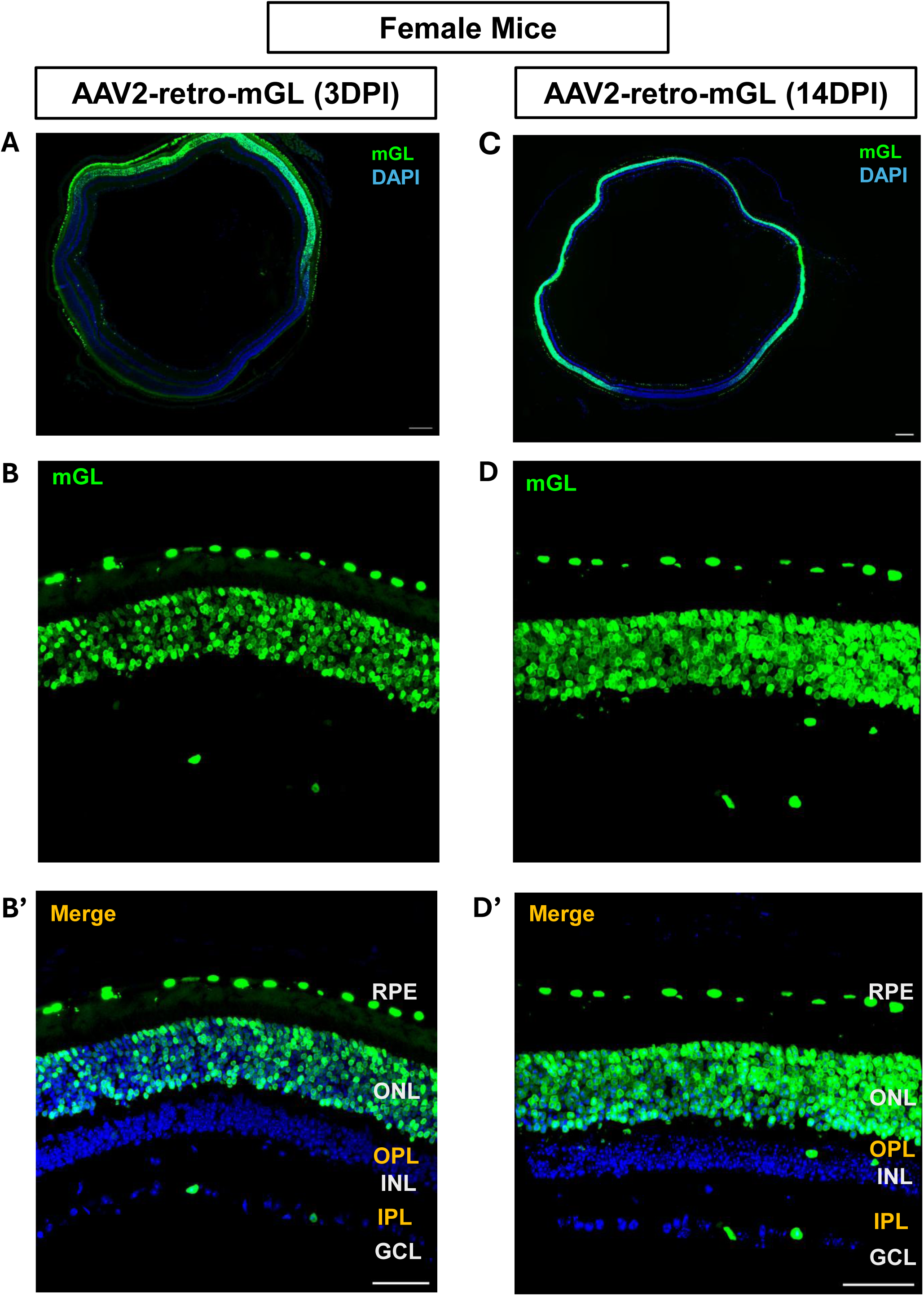
AAV2-retro leads to similar preferential transduction in female mice after intravitreal injection. Stitched 10X images of retinal cross-sections showing mGL expression at 3 dpi **(A)** and 14dpi **(C)**. DAPI (blue) labels nuclei; green indicates mGL reporter expression. **(B and D)** Higher-magnification images of representative retinal regions showing mGL expression at 3 dpi (B) and 14 dpi (D). (B’ and D’) Merged images corresponding to panels B and D, illustrating mGL and DAPI expression across retinal layers. Retinal layers are indicated: RPE, ONL, OPL, INL, IPL, and GCL. Three female mice were examined per time point with similar results. *Scale bars*: 200 µm (A, C); 50 µm (B’, D’).

To rule out the possibility that the observed outer layer specificity resulted from unintended deep injections rather than true intravitreal delivery, we administered wild-type AAV2-mGL via intravitreal injection. This vector is well established to transduce cells primarily in the ganglion cell layer under these conditions. Three days after intravitreal AAV2-mGL injection, mGL expression was detected primarily in the GCL, with no labelling detected in the inner or outer nuclear layer (Fig. 4A). By day 14, transduction was markedly enhanced in the GCL, with dense and specific labeling of ganglion cells; expression remained absent in the outer retina, indicating that AAV2-mGL selectively and efficiently targets ganglion cells over time but do not reach the photoreceptors. (Fig. 4A).

**Figure 4.**
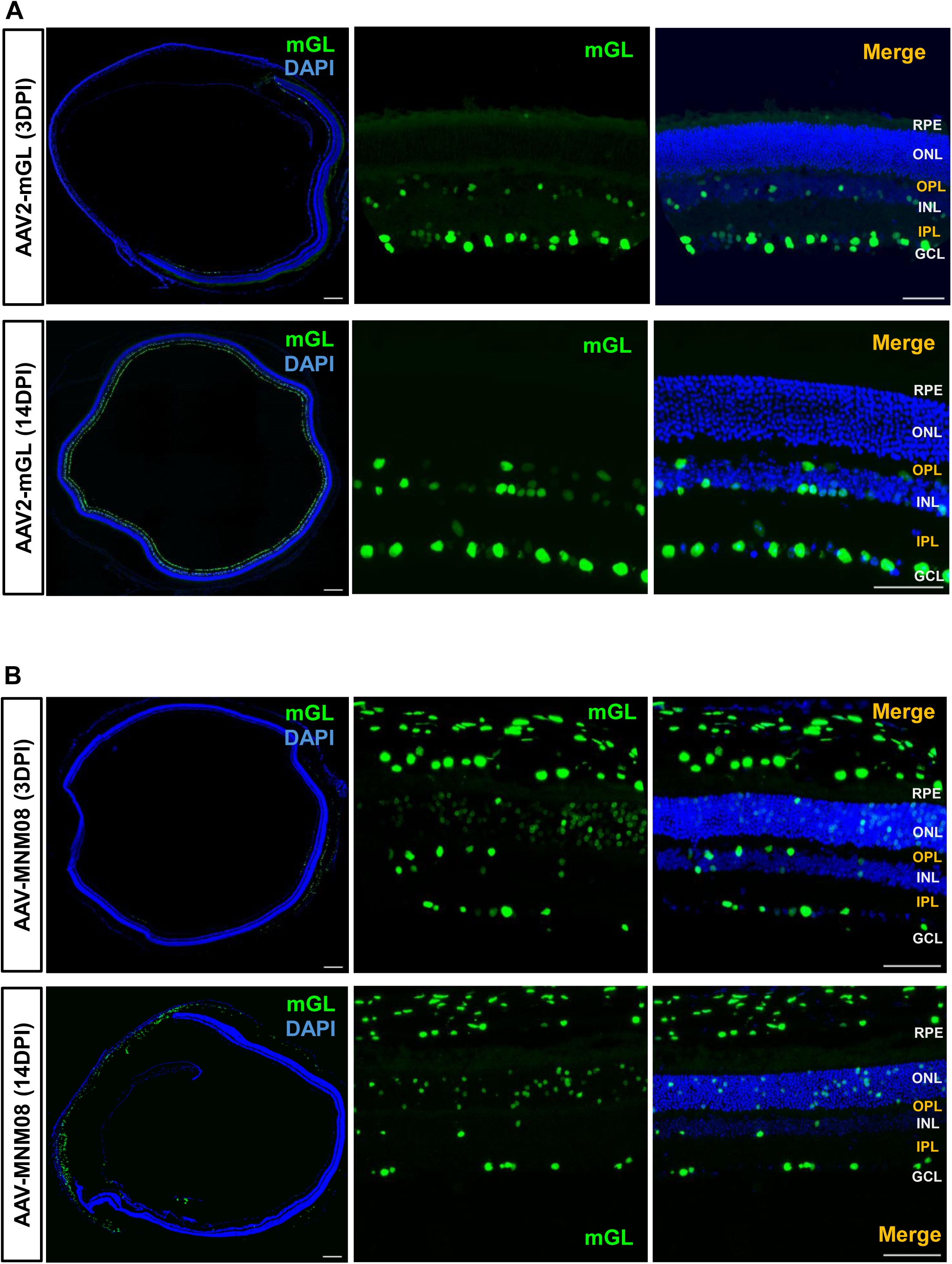
Distinct tropism is seen after intravitreal injection of AAV2-CMV-mGL and AAV2-MNM08-CMV-mGL. **(A)** Retinal cross-sections showing mGL expression at 3 dpi (top row) and 14 dpi (bottom row) following AAV2-CMV-mGL delivery. DAPI (blue) labels nuclei, and green marks mGL reporter expression. **(B)** Retinal cross-sections showing mGL expression at 3 dpi (top row) and 14 dpi (bottom row) following AAV2-MNM08-CMV-mGL delivery. The far-left panels show stitched 10X overviews. Retinal layers are indicated: RPE, ONL, OPL, INL, IPL, and GCL. The total number of animals examined was as follows: AAV2-mGL (3 dpi, n = 5; 14 dpi, n = 5) and AAV-MNM08 (3 dpi, n = 3; 14 dpi, n = 3). *Scale bars*: 200 µm (left panels); 50 µm (right panels).

### Another known retrograde AAV lacks specificity for the outer retinal layers

In addition to the original AAV2-retro^28^, other engineered AAV capsids have been identified that exhibit efficient retrograde transport. Among these, MNM008 capsid, derived from canine adenovirus domains, showed exceptional retrograde infectivity for multiple neuronal types.^11^ To determine whether outer layer–specific gene delivery is a common feature among retrograde AAVs, we compared AAV-MNM008 with AAV2-retro following intravitreal injection. Mice received AAV-MNM008 expressing the mGL reporter, and retinal tissues were analyzed at 3- and 14-days post-injection. We observed that intravitreal injection of AAV-MNM008 results in a distinct transduction profile compared to that of AAV2-retro. At day 3 post injection (Fig. 4B), mGL expression was detected in the photoreceptor layer, but with a lower signal intensity than what we observed with AAV2-retro. At 14dpi, AAV-MNM008 mGL expression was persistent in the GCL and extended to the photoreceptor layer (PRL), although only a limited number of photoreceptor cells were transduced. (Fig. 4B). Together, these findings confirm that AAV2 robustly transduces the RGCs without photoreceptors whereas AAV2-MNM08 enables modest photoreceptor transduction with restricted efficiency and a limited spread. Neither vector matched the outer layer specific transduction profile of AAV2-retro.

### Identification of the retinal cell types transduced by the vectors using cell markers

To identify the retinal cell types transduced by AAV2-retro, we first performed immunostaining with an antibody against cone arrestin (a marker for cone photoreceptors). At 3 and 14dpi, mGL expression colocalized with cone arrestin immunoreactivity, confirming efficient transduction of cones (Figs. 5A, 5B). In addition, mGL^+^ cells lacking cone arrestin immunoreactivity were also observed in the ONL, reflecting rod photoreceptor transduction (Figs. 5A, 5B).

**Figure 5.**
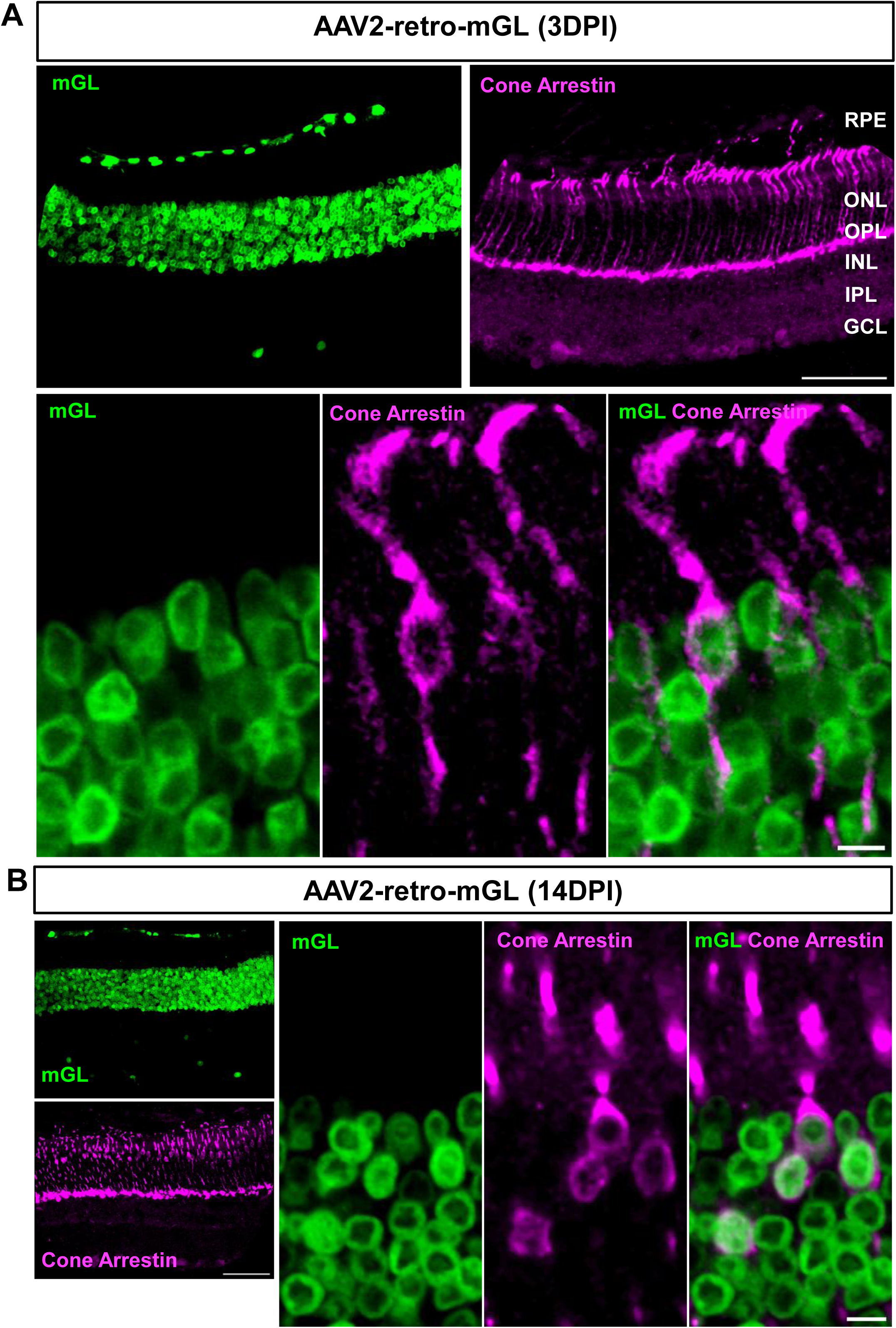
AAV2-retro effectively delivers transgene to both rods and cones. Representative retinal cross-sections showing mGL expression (green) at 3 dpi **(A)** and 14 dpi **(B)** following intravitreal injection of AAV2-retro-CMV-mGL. Cone arrestin (magenta) was used to identify cone photoreceptors. Bottom panel an (A), higher magnification images highlighting the colocalization of mGL with cone arrestin immunoreactivity. *Scale bars*: 50 µm (A, B); 5 µm (higher magnification images).

At both 3 and 14 dpi, a few scattered mGL^+^ cells were also detected in the GCL. To determine the identity of these cells, we performed immunohistochemistry with two cell markers: AP2-α, a marker for amacrine cells and RNA-binding protein with multiple splicing (RBPMS), a marker for RGCs. Staining with AP2-α revealed that most transduced cells in the GCL were amacrine cells. (Figs. 6A, 6B). At 14 days post injection, over 60% of mGL+ cells in the GCL and INL were immunoreactive for AP2-α . In contrast, fewer than 10% of mGL⁺ cells were immunoreactive for RBPMS (Fig. 6C). These results suggest that in the inner retinal layers, where overall transduction was limited, AAV2-retro preferentially targeted amacrine cells, and this tropism persisted over time.

**Figure 6.**
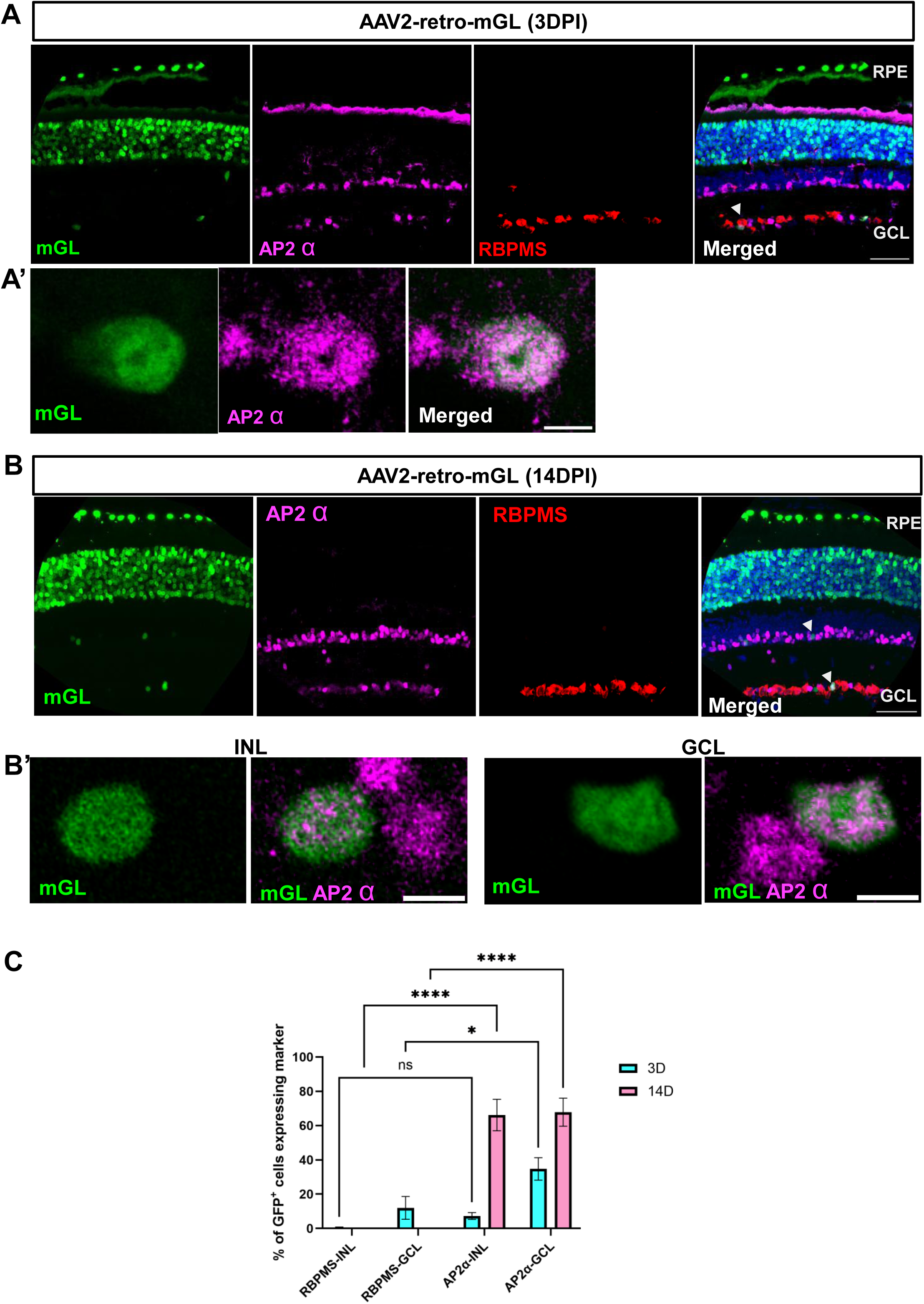
AAV2-retro transduces only a small number of amacrine cells and RGCs. Retinal cross-sections show mGL expression (green) at 3 dpi **(A)** and 14 dpi **(B).** AP-2α immunoreactivity shown in magenta and RBPMS immunoreactivity in red. (A’ and B’) Higher-magnification images showing cells which appear to be co-labelled with mGL and AP-2α. INL, inner nuclear layer; GCL, ganglion cell layer. **(C)** Quantification shows that the percentage of mGL^+^ cells that are positive for either RBPMS (in GCL and INL) or AP-2α at each time point. N=3 per group. Error bar, ± SEM Two-way ANOVA with Tukey’s multiple comparisons: * < 0.05, **** < 0.0001, ns = not significant. *Scale bars*: 50 µm (A, B); 5 µm (A’, B’).

### AAV2-retro transduces cells in the RPE and outer neuroblastic layer of postnatal mouse retina

To assess whether AAV2-retro can transduce retinal cells during early postnatal development, intravitreal injections were performed at postnatal three day (P3) old mice, and retinas were analyzed three days later. Reporter expression was detected predominantly within the RPE and the outer neuroblastic layer (oNBL) (Figure 7). At this developmental stage, rods and cones are already present, however, their maturation is not yet complete, and they progressively contribute to formation of the ONL. Immunostaining with a cone arrestin antibody shows overlap with reporter expressing-cells within the outer oNBL^32,33^, indicating transduction of cone lineage cells during postnatal development (Fig. 7).

**Figure 7.**
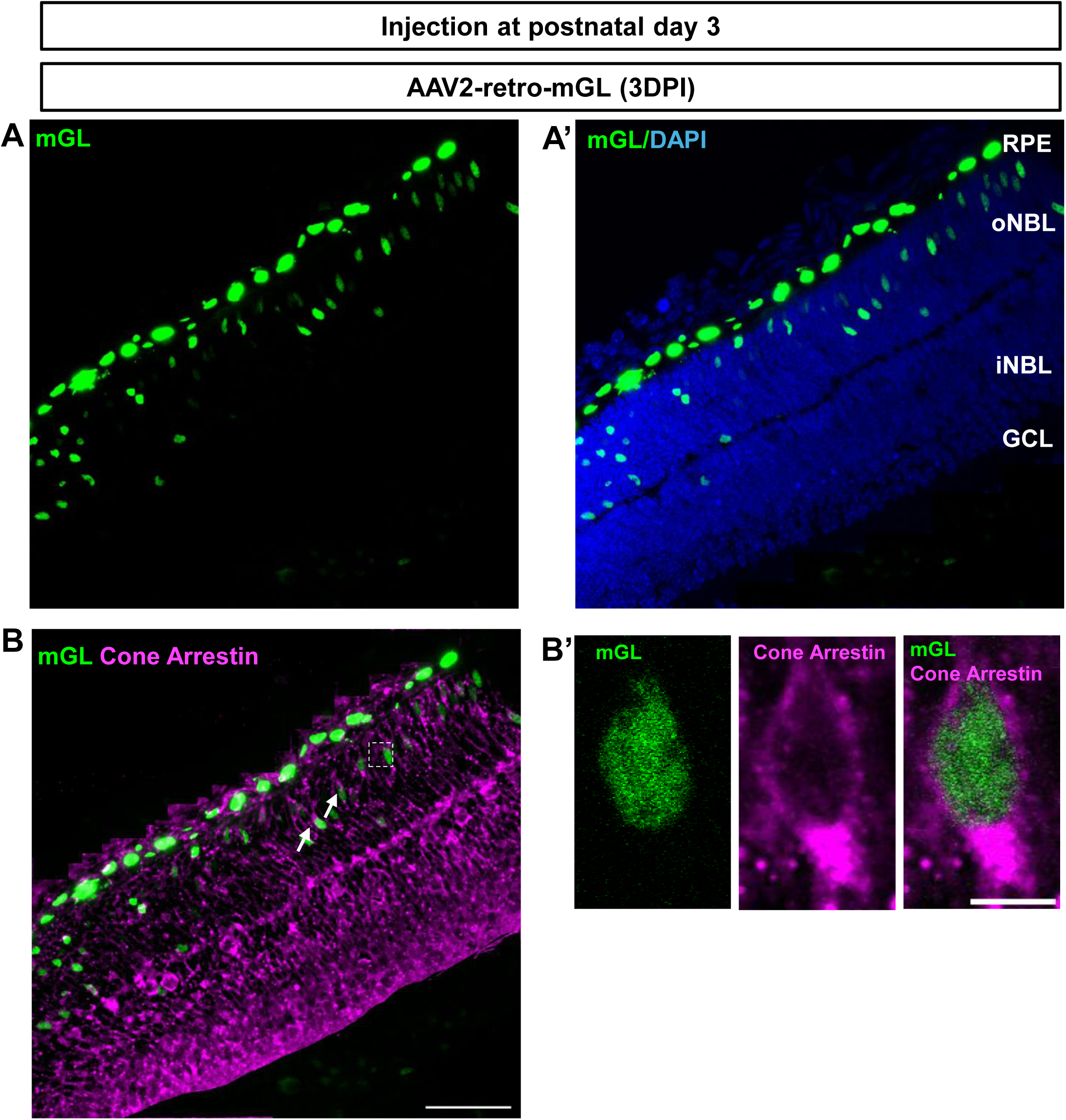
Early postnatal intravitreal delivery of AAV-retro results in viral transduction of RPE cells and cells within the outer neuroblastic layer of the retina. (**A**) Higher-magnification view of a retinal region from a P3-injected pup, showing mGL-positive cells expressed in the outer neuroblastic layer (NBL), consistent with photoreceptors localization. (**A’**) Higher magnification view of the same region showing mGL and DAPI. (B) Immunostaining for cone arrestin (magenta) shows colocalization of mGL and arrestin signal, indicating early cone transduction at when the AAV was injected at P3. (B’) Higher-magnification images highlight cells co-labeled with mGL and cone arrestin. Retinal layers are indicated RPE, oNBL, iNBL, and GCL. AAV was injected intravitreally in four mice with similar results. *Scale bars*: 50 µm (B); 5 µm (B’).

## DISCUSSION

Our results demonstrate that AAV2-retro rapidly and robustly transduces the outer retina, with reporter expression detectable as early as 1 dpi and progressively increasing by 3 and 14 days. We also observed transduction of a few cells in the GCL, where the majority co-express the AP-2α, consistent with the identity of amacrine cells. The AAV2-retro’s pronounced specificity for the outer retinal layers is striking, given that intravitreal delivery has traditionally been considered inadequate for targeting these regions, with most vectors preferentially transducing cells in the INL and GCL.^34,35^ Other engineered AAV capsids, including AAV2.7m8, AAV2.GL, and AAV2.NN have demonstrated markedly improved retinal transduction efficiency compared with wild-type AAV2.^24,36^ However, these variants generally enhance transduction across multiple retinal layers rather than conferring specificity to the outer retina. Consequently, photoreceptor-restricted gene expression in these systems has relied on the use of rod-specific promoters, such as the RHO promoter.^24^ In contrast, the ability of AAV2-retro to overcome these limitations underscores its unique efficiency and distinguishes it from other tested capsid. The absence of a similar expression pattern in MNM08 further underscores the unique characteristics of AAV2-retro.

The specificity and rapid onset of AAV2-retro provide distinct advantages for preclinical animal studies of retinal degeneration. When delivered intravitreally, AAV2-retro efficiently and selectively transduces photoreceptors and RPE within a brief period, eliminating the need for invasive subretinal injection. This property allows for more consistent and reproducible gene delivery across the retina and reduces surgical variability and tissue damage. The rapid onset of transgene expression further facilitates early evaluation of gene function and therapeutic efficacy, which is a key advantage in disease models characterized by fast progressing degeneration. Taken together, these attributes make AAV2-retro a potent tool for accelerating preclinical testing of gene therapies targeting photoreceptor and RPE dysfunction. Importantly, confining gene manipulation to photoreceptors and RPE minimizes off-target effects from transgene expression in inner retinal neurons or glial cells, which could otherwise perturb retinal circuitry and impair visual function. Gene expression in non-targeted retinal cell types can alter synaptic signaling, disrupt homeostatic support functions, or induce unintended stress responses, ultimately confounding the interpretation of experimental outcomes. By confining transgene delivery to the outer retina, AAV2-retro enables more precise assessment of gene function within disease-relevant cell populations while preserving the physiological integrity of the inner retinal network. Thus, the unique tropism of AAV2-retro raises opportunities for studying outer retinal physiology in vivo. By restricting transgene expression to photoreceptors and RPE, researchers can manipulate signaling pathways, visual cycle components, or metabolic processes without disrupting inner retinal neuronal networks, enabling clearer mechanistic insights.

While single intravitreal injections resulted in strong photoreceptor and RPE transduction, retinal coverage was not uniform; second intravitreal injection further enhanced both the efficiency and spatial coverage of transduction, suggesting that sequential injections can be used to overcome local variability in viral entry and improve gene delivery outcomes.

In this study, we also demonstrate that AAV2-retro exhibits similarly robust tropism for photoreceptors and RPE in female mice, paralleling the strong and selective outer retinal transduction observed in males. The consistency of vector performance across sexes is an important finding, as several retinal degenerative diseases including certain forms of age-related macular degeneration, autoimmune retinopathies, and mitochondrial optic neuropathies show higher prevalence, faster progression, or distinct clinical manifestations in females. The ability of AAV2-retro to reliably target the outer retina in female animals underscores its utility as a gene-delivery tool for modeling and potentially treating these sex-skewed conditions.

We further show that AAV2-retro delivered intravitreally at P3 can efficiently transduce the RPE and the oNBL, including cone lineage cells, during a critical window of early retinal development. Many inherited retinal degenerations such as Leber congenital amaurosis and early-onset retinitis pigmentosa may cause photoreceptor dysfunction or death beginning in early postnatal or even prenatal stages. Thus, the ability of AAV2-retro to access and transduce photoreceptor precursors at P3 suggests that this vector may be particularly valuable in animal models where therapeutic intervention must occur before maturation of the outer nuclear layer. By providing efficient early developmental gene delivery via a simple intravitreal injection, AAV2-retro offers a practical tool for probing disease pathogenesis and evaluating therapeutic rescue in IRD models where early photoreceptor targeting is essential.

The mechanism underlying AAV2-retro’s selective tropism for outer retinal cells remains unclear. One possibility is that its capsid structure facilitates enhanced penetration through the inner limiting membrane, a barrier that has historically limited the efficacy of intravitreal gene transfer.^37^ Alternatively, AAV2-retro may possess capsid motifs that preferentially interact with cell-surface receptors expressed on photoreceptors or RPE, enabling rapid internalization and transgene expression. Given the vector’s original design for retrograde transport in CNS neurons^28,38^ its adaptation to the retinal environment may involve unexpected cross-compatibility with outer retinal cellular entry pathways.

Despite the promise of AAV2-retro, several limitations warrant consideration. Although our results demonstrate rapid and efficient transgene expression, the long-term stability of this transduction remains to be determined. Comprehensive evaluation of expression durability, potential immune responses, and the safety of repeated dosing in larger cohorts may be needed. In a previous study, several AAV capsids, including AAV2-retro, were evaluated for their ability to transduce cells in the human retina ex vivo. In this experiment, human retinal explants were cultured in transwells with the photoreceptor layer facing downward, and AAVs were applied from the vitreal side (inner limiting membrane/retinal ganglion cell layer) to mimic intravitreal injection conditions. Under these conditions, AAV2-retro produced only limited transduction of the human retina.^39^ Although ex vivo and in vivo environments differ substantially, these findings suggest that the robust outer retinal transduction observed in mouse eyes in the present study may not directly translate to other species.

In summary, our work establishes AAV2-retro as a markedly efficient virus for selectively targeting photoreceptors and RPE following intravitreal injection. Its rapid onset of expression, superior efficiency compared with established serotypes, and enhanced distribution with sequential dosing underscore its potential for experimental applications.

