## Supplementary Figure for "AAV2-Retro-Mediated Gene Transfer Selectively Targets Outer Retinal Cells Following Intravitreal Injection"

**Affiliations:**

### Supplementary Figure

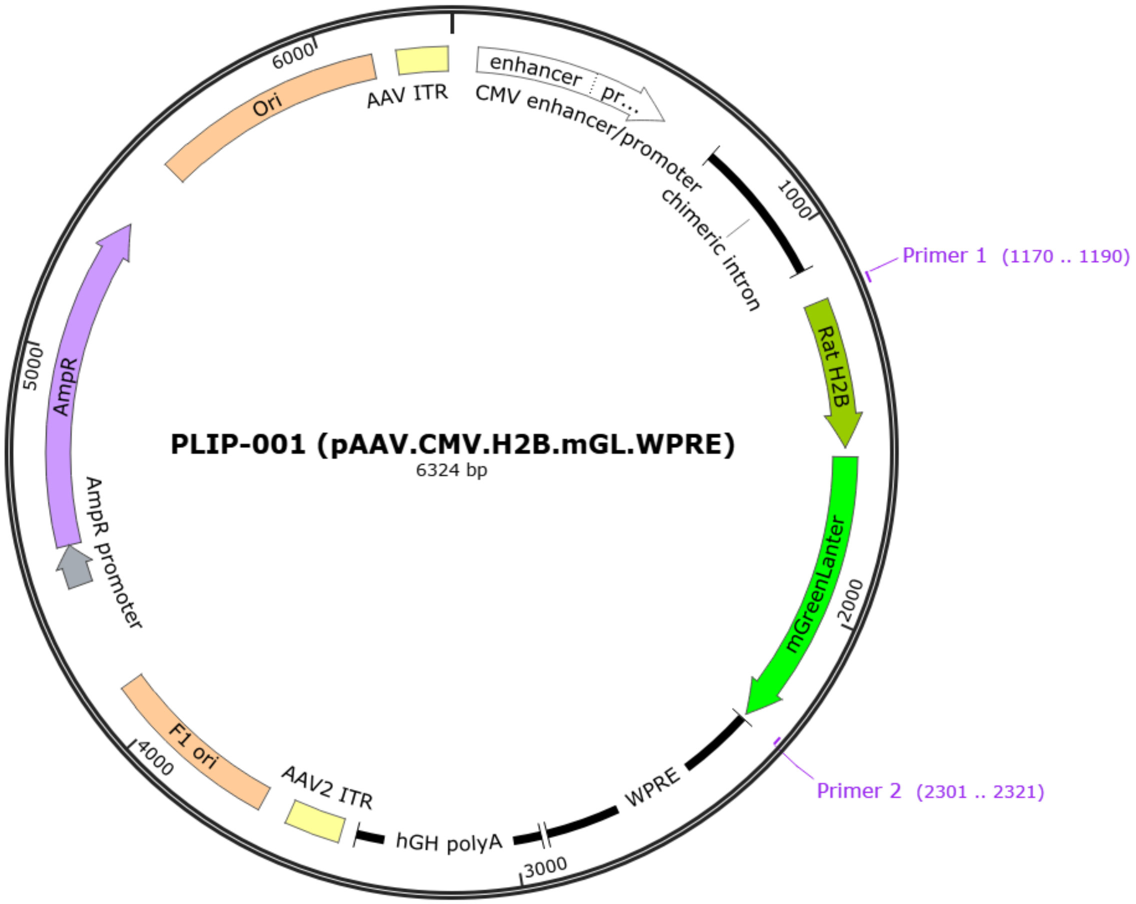


**Figure S1: The Map of AAV2-retro-CMV-mGL vector.** The transgene cassette comprises a CMV promoter driving mGL fused to H2B, followed by WPRE and hGH polyA, flanked by AAV2 ITRs.
